# The genetic architecture of the network underlying flowering time variation in *Arabidopsis thaliana*

**DOI:** 10.1101/175430

**Authors:** Eriko Sasaki, Florian Frommlet, Magnus Nordborg

## Abstract

Flowering time is a key adaptive trait in plants and is tightly controlled by a complex regulatory network that responds to seasonal signals. In a rapidly changing climate, understanding the genetic basis of flowering time variation is important for both agriculture and ecology. Genetic mapping has revealed many genetic variants affecting flowering time, but the effects on the gene regulatory networks in population-scale are still largely unknown. We dissected flowering time networks using multi-layered Swedish population data from *Arabidopsis thaliana*, consisting of flowering time and transcriptome collected under constant 10°C growth temperature in addition to full genome sequence data. Our analysis identified multiple alleles of the key flowering time gene *FLOWERING LOCUS C* (*FLC*) as the primary determinant of the network underlying flowering time variation under our condition. Genetic variation of *FLC* affects multiple-pathways through known flowering-time genes including *FLOWERING LOCUS T* (*FT*), and *SUPPRESSOR OF OVEREXPRESSION OF CONSTANS 1* (*SOC1*). We demonstrated that an extremely simple single-locus model of *FLC* involving allelic variation and expression explains almost a half of flowering time variation, with 60% of the effect being mediated through *FLC* expression. Furthermore, the accuracy of the model fitted at 10°C is almost unchanged at 16°C.

## Introduction

Timing of reproduction is a key adaptive strategy in plants. To decide when to flower, plants integrate a number of seasonal signals like day length, temperature, and humidity (SIMPSON and DEAN 2002; KIM *et al.* 2009; ANDRES and COUPLAND 2012). Understanding the mechanisms controlling flowering time, and the genetic architecture of variation for this trait is essential for agriculture as well as for predicting how plants will respond to climate change. It is also a model for selection on a complex, adaptive trait. The regulation of flowering is one of the best-studied developmental transitions in plants. In *A. thaliana*, a complex network including more than one hundred genes in several major pathways has been described: the photoperiod, ambient temperature, autonomous, integrator, gibberellin and vernalization pathways combine to regulate flowering (SIMPSON and DEAN 2002; KIM *et al.* 2009; WELLMER and RIECHMANN 2010; SRIKANTH and SCHMID 2011; ANDRES and COUPLAND 2012). Many mathematical and statistical models of flowering time regulation have been proposed based on genetic data (WELCH *et al.* 2003; SATAKE and IWASA 2012; SATAKE *et al.* 2013; LI *et al.* 2014b; WANG *et al.* 2014; LEAL VALENTIM *et al.* 2015), as well as time-course data of expression levels of known flowering time genes (SCHMID *et al.* 2003). In contrast, relatively little has been done in terms of modeling the pathways that lead to natural variation for flowering time. SATAKE *et al.* (2013) investigated the dynamics of the vernalization pathway and its variation using two individuals of *A. halleri* using expression levels of marker genes, but variation in the flowering network on a population scale is still poorly understood. In this study, we present a model of flowering time network variation in a population of *A. thaliana*. Our primary goal was to investigate how gene expression data combined with genetic variation data might help us understand the regulatory networks that connect genotypes to phenotype. To build the model, we take advantage of a multi-layered data set of *A. thaliana* from Sweden that contains genotypes (LONG *et al.* 2013), RNA-seq transcriptome data (DUBIN *et al.* 2015), as well as flowering time phenotypes (SASAKI *et al.* 2015) for 132 individuals.

## Results

### The correlation between gene expression and flowering time

We began by asking whether gene expression, as measured in whole plants (above ground only) at a few weeks of age (the nine-leaf stage) was correlated with eventual flowering of the same genotype (in 10°C, long day conditions) across 132 inbred lines (Table S1). Using a significance threshold corresponding to an FDR of 5%, 38 out of 20,285 genes (0.2%) showed significant correlation with flowering time (see Methods and Table 1). Of these, 9 were annotated as being related to a flowering time phenotypes, and 5 were also part of a more conservative list of *a priori* candidates (SRIKANTH and SCHMID 2011). This represents a highly significant enrichment, which persists at higher FDR cut-offs (Figure 1A). The correlations also generally remain after a standard correction for population structure using a linear mixed model (Table 1; see Methods).

**Table 1.** List of genes whose expression is significantly correlated with flowering time (10°C)

| Gene ID | $\rho$ | $r^2$ | $p$-value (LM) | $p$-value (LMM) | Description <sup>a</sup> |
| --- | --- | --- | --- | --- | --- |
| AT5G10140 | 0.63 | 0.40 | 3.05E-16 | 9.30E-11 | **FLC\*** |
| AT1G65480 | -0.54 | 0.29 | 2.64E-11 | 3.32E-08 | **FT\*** |
| AT2G45660 | -0.47 | 0.22 | 1.35E-08 | 5.21E-05 | **SOC1\*** |
| AT2G41640 | -0.42 | 0.17 | 7.03E-07 | 4.30E-05 | Glycosyltransferase |
| AT3G57920 | -0.39 | 0.15 | 3.28E-06 | 2.58E-02 | **SPL15** |
| AT1G04400 | -0.38 | 0.15 | 5.24E-06 | 1.39E-02 | **CRY2\*** |
| AT5G52310 | -0.38 | 0.15 | 5.39E-06 | 6.72E-04 | *RD29A* |
| AT1G69440 | -0.38 | 0.15 | 5.53E-06 | 2.82E-03 | *AGO7* |
| AT3G13100 | -0.38 | 0.14 | 7.71E-06 | 1.99E-04 | ATP-BINDING CASSETTE C7 |
| AT1G23870 | -0.38 | 0.14 | 8.98E-06 | 2.20E-03 | *TPS9* |
| AT5G44630 | -0.37 | 0.14 | 9.65E-06 | 2.83E-04 | Terpenoid cyclases |
| AT3G09100 | -0.37 | 0.14 | 9.74E-06 | 3.15E-03 | protein coding |
| AT5G51720 | 0.37 | 0.14 | 9.90E-06 | 2.11E-01 | **AT-NEET** |
| AT4G33040 | -0.37 | 0.14 | 1.02E-05 | 1.14E-02 | protein coding |
| AT3G04485 | 0.37 | 0.13 | 1.51E-05 | 9.65E-03 | other RNA |
| AT1G77810 | -0.37 | 0.13 | 1.62E-05 | 4.17E-04 | Galactosyltransferase |
| AT2G13560 | -0.36 | 0.13 | 1.70E-05 | 2.69E-02 | *NAD-ME1* |
| AT3G08990 | 0.36 | 0.13 | 1.73E-05 | 6.09E-02 | protein coding |
| AT1G17020 | -0.36 | 0.13 | 1.78E-05 | 2.24E-04 | *SRG1* |
| AT1G06160 | 0.36 | 0.13 | 2.26E-05 | 4.11E-02 | *ORA59* |
| AT3G19860 | -0.36 | 0.13 | 2.35E-05 | 5.81E-04 | *BHLH121* |
| AT5G48400 | -0.36 | 0.13 | 2.60E-05 | 6.83E-04 | *ATGLR1.2* |
| AT3G19500 | 0.36 | 0.13 | 2.76E-05 | 6.12E-04 | protein coding |
| AT3G05660 | -0.36 | 0.13 | 2.80E-05 | 5.79E-02 | *AtRLP33* |
| AT4G24540 | -0.35 | 0.12 | 3.33E-05 | 1.38E-02 | **AGL24\*** |
| AT5G25120 | -0.35 | 0.12 | 3.42E-05 | 5.08E-03 | *CYP71B11* |
| AT3G18840 | -0.35 | 0.12 | 4.03E-05 | 9.30E-03 | TPR-like superfamily protein |
| AT2G18196 | 0.35 | 0.12 | 4.67E-05 | 2.06E-03 | protein coding |
| AT5G46210 | -0.35 | 0.12 | 4.78E-05 | 2.21E-03 | **ATCUL4** |
| AT1G53165 | -0.35 | 0.12 | 5.01E-05 | 2.32E-04 | ATMAP4K ALPHA1 |
| AT3G20250 | -0.34 | 0.12 | 5.12E-05 | 1.19E-04 | *APUM5* |
| AT5G44590 | 0.34 | 0.12 | 5.68E-05 | 2.67E-02 | protein coding |
| AT3G55610 | -0.34 | 0.12 | 6.47E-05 | 2.36E-04 | *P5CS2* |
| AT4G18130 | -0.34 | 0.12 | 6.63E-05 | 6.13E-04 | **PHYE** |
| AT1G78050 | -0.34 | 0.12 | 6.82E-05 | 1.22E-03 | *PGM* |
| AT5G10490 | -0.34 | 0.12 | 6.94E-05 | 4.42E-04 | *MSL2* |
| AT5G58900 | 0.34 | 0.11 | 7.22E-05 | 1.37E-01 | protein coding |
| AT2G46500 | -0.34 | 0.11 | 7.92E-05 | 7.57E-04 | *ATPI4K* |

<sup>a</sup> Genes in bold have flowering-related mutant phenotypes; \*denotes genes that are also part of a more conservative list of *a priori* candidate (SRIKANTH and SCHMID 2011).

**Figure 1.**
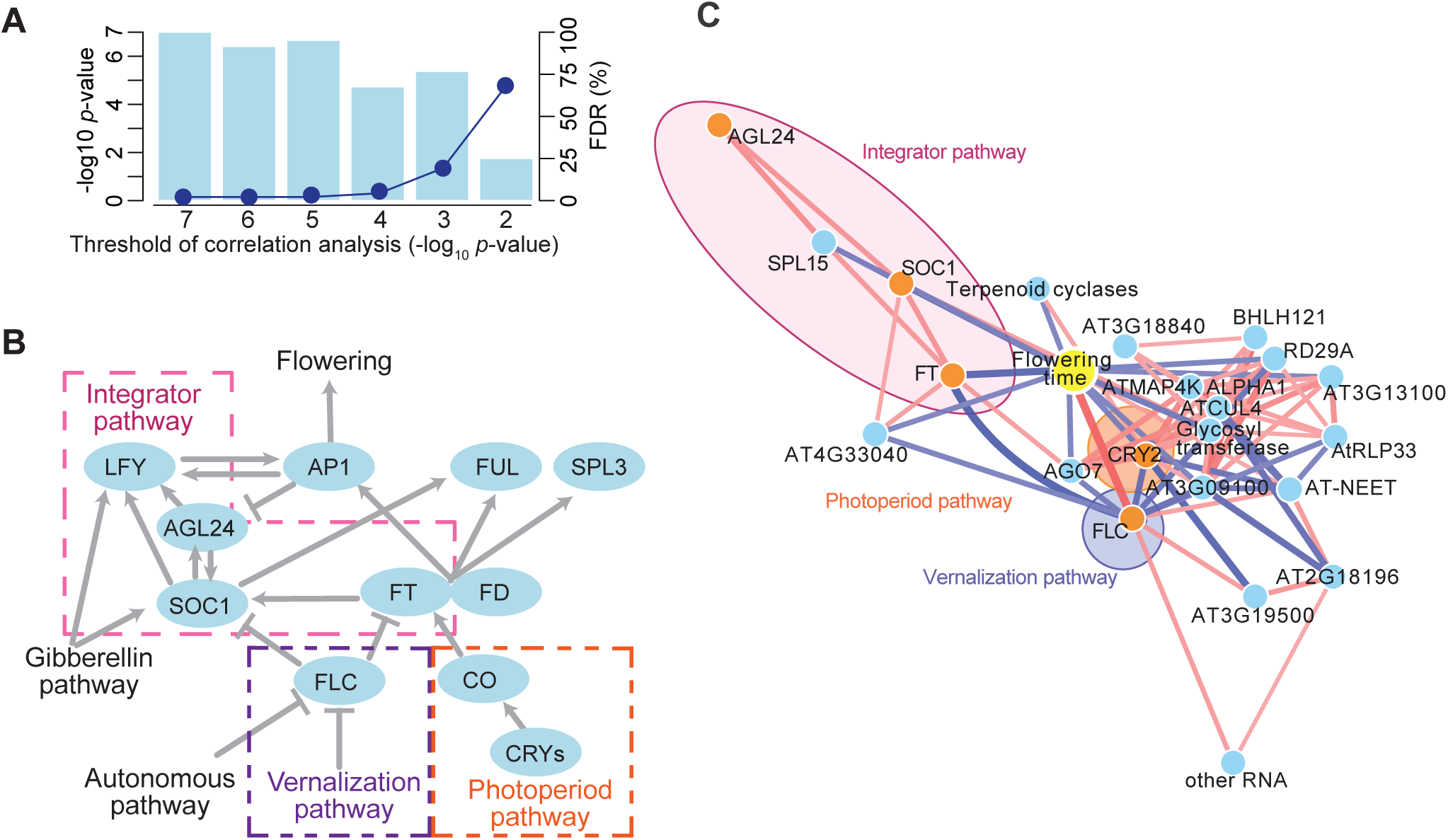
Correlation between flowering time and gene expression levels in the Swedish population. (A) The significance of the GO enrichment for flowering time genes (and implied FDR; see Methods) as function of the significance threshold for the flowering-expression correlation. (B) Outline of the flowering pathways in *A. thaliana* (reviewed in, *e.g.*, KIM *et al.* 2009; WELLMER and RIECH-MANN 2010; SRIKANTH and SCHMID 2011). FLC represses the floral integrator genes FD, FT, and SOC1. FT is induced by the photoperiod pathway through CONSTANS (CO), which is induced by CRYPTOCROMEs (CRYs); the FT protein is a mobile flowering signal that works with FD to induce SOC1 and floral meristem genes including APETALA1 (AP1), FRUITFUL (FUL), and SEPALATA (SPL3). AGL24 and SOC1 regulate each other in positive feedback loops and induce transcription of LFY. The gibberellin pathway promotes flowering by inducing SOC1 and the floral meristem-identity gene LEAFY (LFY). (C) A correlation network based on measured expression level. Nodes show flowering time (yellow) and the genes in Table 1 (blue, or orange for the *a priori* gene set). Edges show significant correlations between nodes (*p*-value < 0.01 with Bonferroni correction) in pink or blue (for positive and negative correlations, respectively).

The top three genes (Table 1) were all *a priori* flowering time genes: *FLOWERING LOCUS C* (*FLC*; MICHAELS and AMASINO 1999; SHELDON *et al.* 1999) in the vernalization pathway, *FLOWERING LOCUS T* (*FT*; KARDAILSKY *et al.* 1999; KOBAYASHI *et al.* 1999) and *SUPPRESSOR OF OVEREXPRESSION OF CONSTANS 1* (*SOC1*; SAMACH *et al.* 2000) in the “integrator” pathway (Table 1; Figure 1B). In agreement with previous work, *FLC* expression was clearly most strongly correlated: the explained variance, *r*^2^ = 0.40, is strikingly similar to what was seen by LEMPE *et al.* (2005) using a different sample under environmental conditions. The expression of the integrator loci *FT* and *SOC1* is less strongly correlated with flowering, which is interesting given that these loci are supposed to act downstream of *FLC*, and are in this sense closer to the phenotype (Figure 1B; SCHMID *et al.* 2003; WELLMER and RIECHMANN 2010).

The correlation network connecting the genes in Table 1 with flowering (see Methods) was consistent with the known flowering-time pathways (Figure 1C). The integrator pathway connected *FT* and *SOC1* with another strong *a priori* candidate, *AGL24*, a known inducer of *SOC1* (YU *et al.* 2002, 2004; MICHAELS *et al.* 2003). The photoperiod pathway was not connected with the integrator pathway, but included *CRY2* (TOTH *et al.* 2001) as a hub gene in a network containing 19 other genes. The vernalization pathway, via *FLC*, cleary plays a central role, connecting the integrator pathway and the photoperiod pathways via *FT* and *CRY2*.

### The genetic basis of flowering-associated expression variation

A network based on expression correlation is inherently undirected and tells us little about causation, however some insight can be gained by identifying the genetic causes of the expression variation (SCHADT *et al.* 2005). We used variance component analysis (LIPPERT *et al.* 2014; MENG *et al.* 2016) to estimate the effect on gene expression of the local genetic variation using a 30 kb window surrounding each gene. Based on permutation tests (*p*-value < 0.05), almost one third of the genes in Table 1 had the property that genetic variation surrounding the gene contributed significantly to the expression of that gene (i.e., they are *cis*-regulated; see Figure 2 and Table S2). *FLC* stood out in that not only was it strongly *cis*-regulated, but genetic variation at the gene was also strongly associated with half of the other genes in Table 1 (Figure 2; Table S2). Thus genetic variation at *FLC* is causing the expression variation at these other loci, almost certainly through its effect on *FLC* expression. In contrast, the expression level of several genes highly correlated with flowering time, including *FT*, *SOC1*, and *CRY2* showed no evidence of *cis*-regulation, but strong evidence for being regulated by genetic variation at *FLC*. This result suggests that *FLC* is the key determinant of flowering time under our conditions.

**Figure 2.**
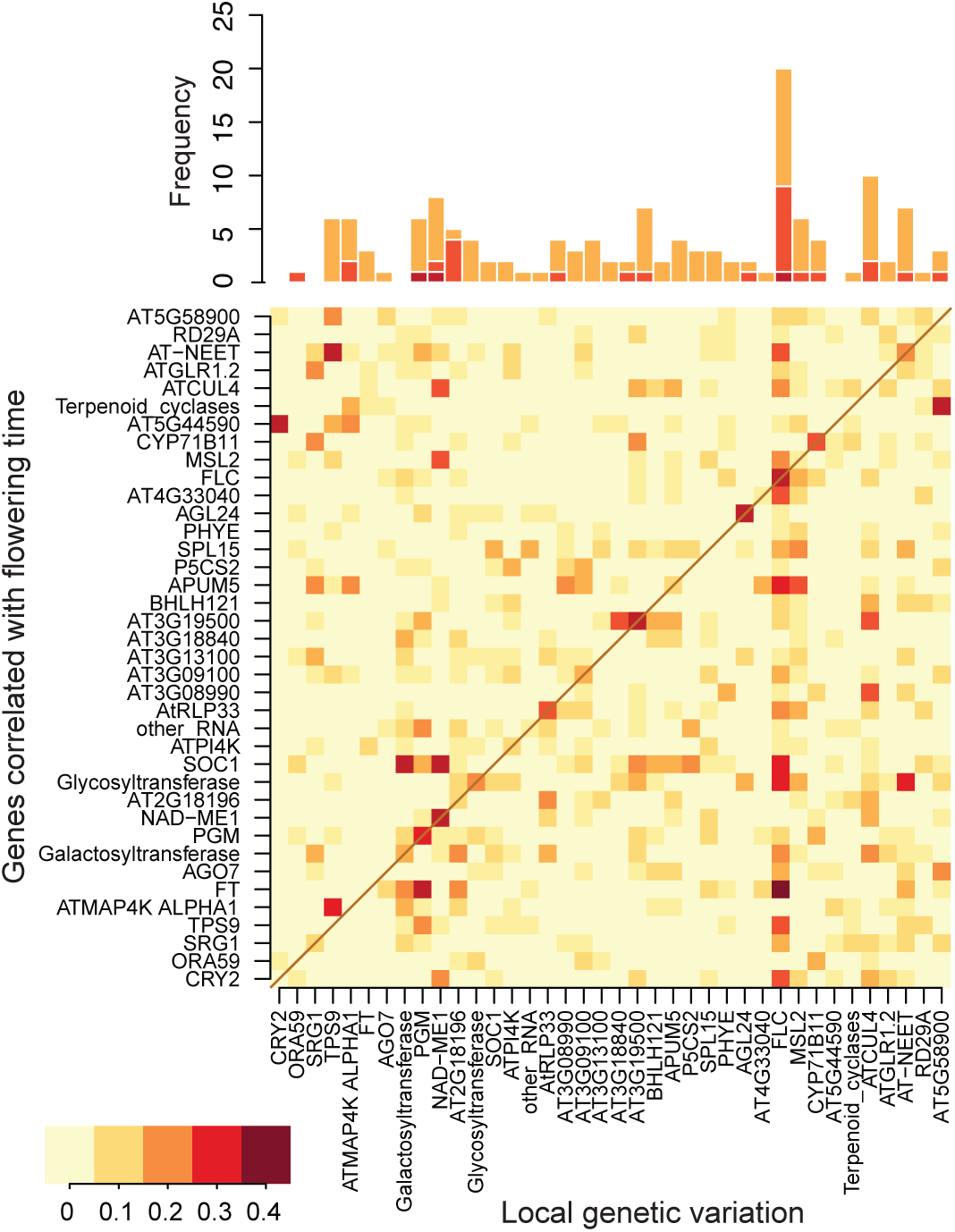
Genetic effects on gene expression levels. Effects of local genetic variation were estimated using a variance component analysis and 30 kb windows surrounding each gene. The bottom panel shows fraction of the gene expression variation (for each gene in Table 1) explained by local genetic variation surrounding each gene in Table 1; the top panel shows frequency of strong associations (*≥*10%). The diagonal line indicates the *cis*-regulation for each gene.

### The genetic basis of flowering time and FLC expression variation

To gain further insight into the contribution of *FLC* to flowering time variation, we carried out genome-wide association studies (GWAS) for flowering time and *FLC* expression (Figure 3, S1). In agreement with our previous results (SASAKI *et al.* 2015), GWAS for flowering time identified a genome-wide significant association with a single nucleotide polymorphism (SNP) in the promoter region of *FLC* (Chr5: 3,180,721; *p*-value = 1.14E-08, MAF = 0.62) in addition to weaker associations in two other *a priori* candidates (Figure 3A). On the other hand, GWAS for *FLC* expression did not identify any significant association (Figure 3B), even within the *FLC* locus itself—which is surprising given the strong correlation with flowering time (Figure 3C) and the evidence for *cis*-regulation obtained using variance-components analysis (Figure 2).

**Figure 3.**
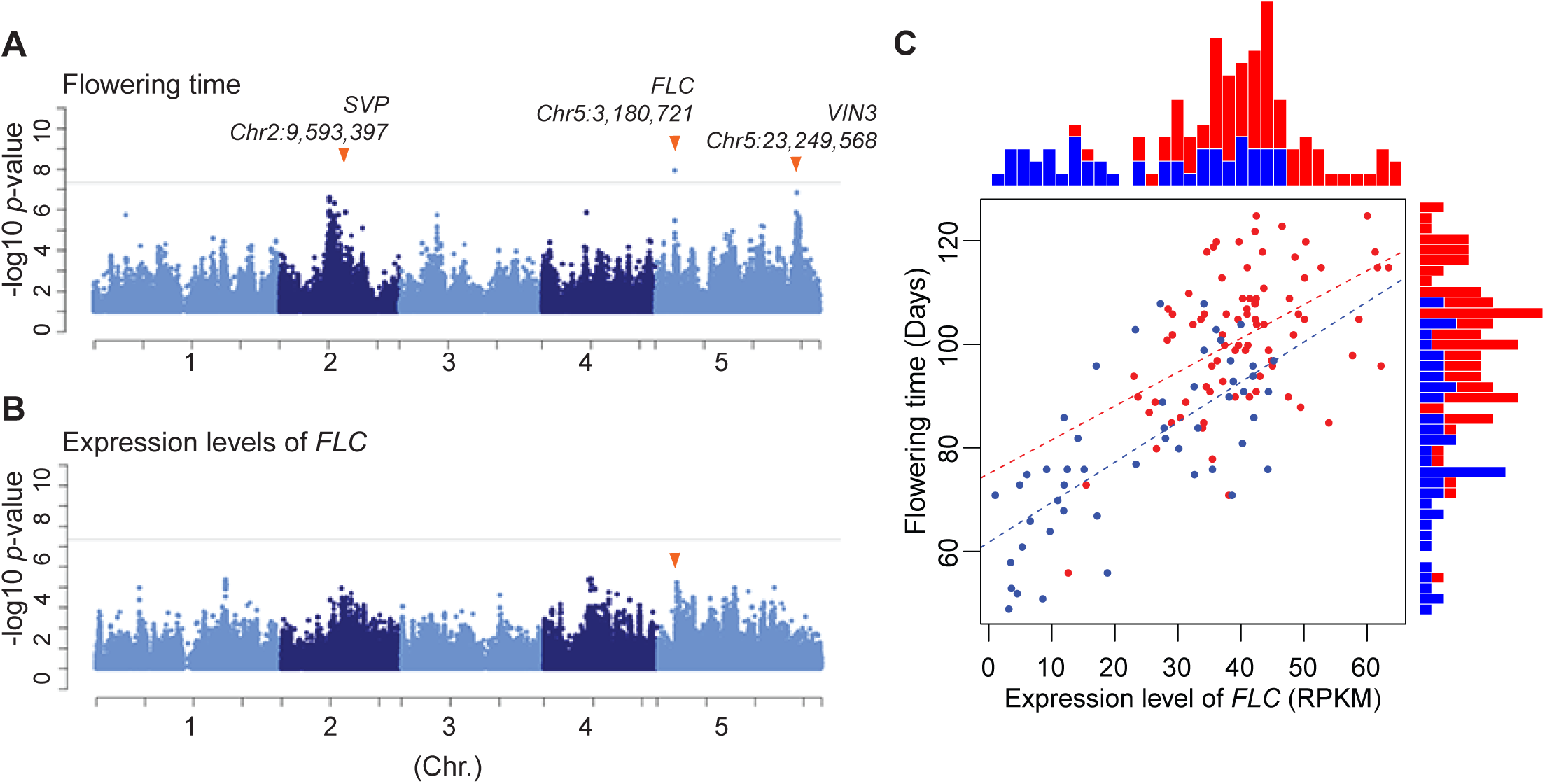
GWAS for flowering time (A) and the *FLC* expression (B). Gray horizontal lines indicate Bonferroni-correct 5% significance thresholds and orange arrows in panel A show *a priori* flowering time genes (from SASAKI *et al.* (2015); the arrow in B shows the SNP in the *FLC* region identified in A. (C) A scatter plot between flowering time and the expression level of *FLC*, and histograms of the genotype in SNP_*FLC*_. Reference and non-reference alleles are shown in blue and red, respectively. Dot lines are regression lines for each allele.

### The genetic architecture of flowering time variation

We are thus faced with a seemingly paradoxical result. How can a SNP at *FLC* (SNP_*FLC*_) predict flowering time but not *FLC* expression, when *FLC* expression strongly predicts flowering time (Figure 3C)? A simple answer would be variation at the protein level, but there is no non-synonymous variation in this gene (LI *et al.* 2014a), and indeed the variance component analysis confirms that the genetic variation is *cis*-regulatory (Figure 2).

The obvious conclusion is that SNP_*FLC*_ must be associated with some aspect of *FLC* expression that is not captured by our expression data, and that the expression variation we measure must be partly caused by *FLC* variation not tagged by SNP_*FLC*_ (in addition to *trans*-acting genetic variation). The variance-components analysis supports the latter explanation: To gain insight into the former, we resorted to a statistical mediation analysis (BARON 1986; VALERI and VANDERWEELE 2013; PALMER *et al.* 2017). A mediation analysis is a model-based attempt to dissect mechanisms underlying an observed relationship between a factor (**A**: exposure), an outcome (**Y**), and an intermediate factor (**M**: mediator). The total effect of **A** on **Y** is decomposed into an indirect effect mediated by **M** and a residual direct effect. In the present context, we assumed that the SNP_*FLC*_ (**A**) regulates flowering time (**Y**) and that this effect is partly mediated through the measured expression level (**M**). To consider the effect of population structure on both **M** and **Y**, we implemented a linear mixed model that took genetic background into account instead of using a standard generalized linear model (Figure 4A; see also Methods and Supplemental Note).

**Figure 4.**
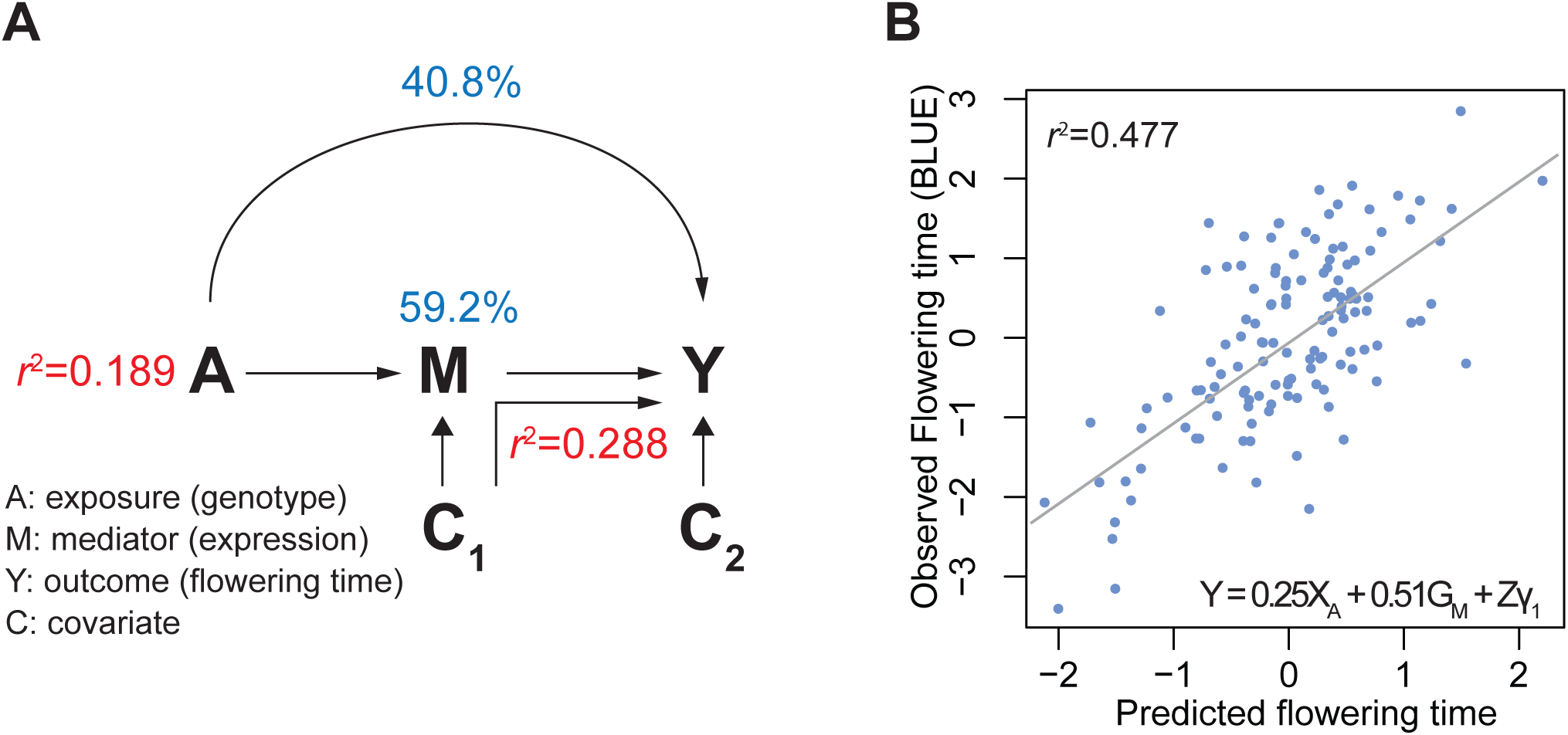
Network structure of flowering time regulation by *FLC*. (A) A mediation model of the flowering time regulation under the control of SNP_*FLC*_. Flowering time variation explained by total SNP_*FLC*_ and direct *FLC* expression level are shown in red, direct and indirect effect size of SNP_*FLC*_ are shown in blue. (B) Predicted flowering time by a FLC_10*°* *C*_ full model. **X** is genotype of SNP_*FLC*_, **G** is *FLC* expression, **Z** is polygenic effects, and *γ*_1_ is a random effect corresponding to the genetic background.

SNP_*FLC*_ explained 19% of flowering time variation in our GWAS. According to the mediation model, only 59% of this effect is mediated by the measured *FLC* expression level, with the remaining 41% being the direct effect — which, per the argument given above, must correspond to unmeasured effects on *FLC* regulation. In addition, *FLC* expression levels also affected flowering independently of SNP_*FLC*_, presumably due to a combination of *cis*-acting variation not captured by SNP_*FLC*_ and *trans*-acting genetic background effects not captured by the kinship matrix. This effect explained 29% of flowering time variation. In total, the full model including SNP_*FLC*_ and *FLC* expression explained a massive 48% of flowering time (Figure 4B).

### Prediction of flowering time using the FLC model

To investigate the limits of our model for prediction, we first tested our model on flowering time and expression data generated for the same population, but at a higher growth temperature that prevent vernalization (DUNCAN *et al.* 2015), namely 16*°*C (DUBIN *et al.* 2015; SASAKI *et al.* 2015). We predicted flowering time using the FLC_10*°* *C*_ model with parameters estimated using the 10*°*C data (Figure 4B). The effect of population structure was estimated using the 16*°*C *FLC* expression levels (see Methods).

SNP_*FLC*_ was significantly associated with flowering time in these data as well (*p*-value = 3.31E-07; MAF = 0.72; Figure S2A-B), but the global correlation of *FLC* expression with flowering time decreased from *R* = 0.63 (at 10*°*C; Table 1) to *R*=0.47 (*p*-value=4.76E-12; Table S3). The correlation was observed in only early flowering lines. Regardless of this, the efficiency of the FLC_10_*°* _*C*_ model changed surprisingly little, and 43% of flowering time variation was predicted by the model (Figure 5A-B, E). We also tested the model on a different population for which flowering data (at around 23*°*C in a greenhouse) and *FLC* expression data were available. In these data SNP_*FLC*_ was not significantly associated with flowering time, suggesting that that *trans*-acting loci break the correlation under higher growth temperature (Figure S2C-D). However, *FLC* expression still showed a weak correlation with flowering, and the model predicted 29% of flowering time variation (Figure 5C-E).

**Figure 5.**
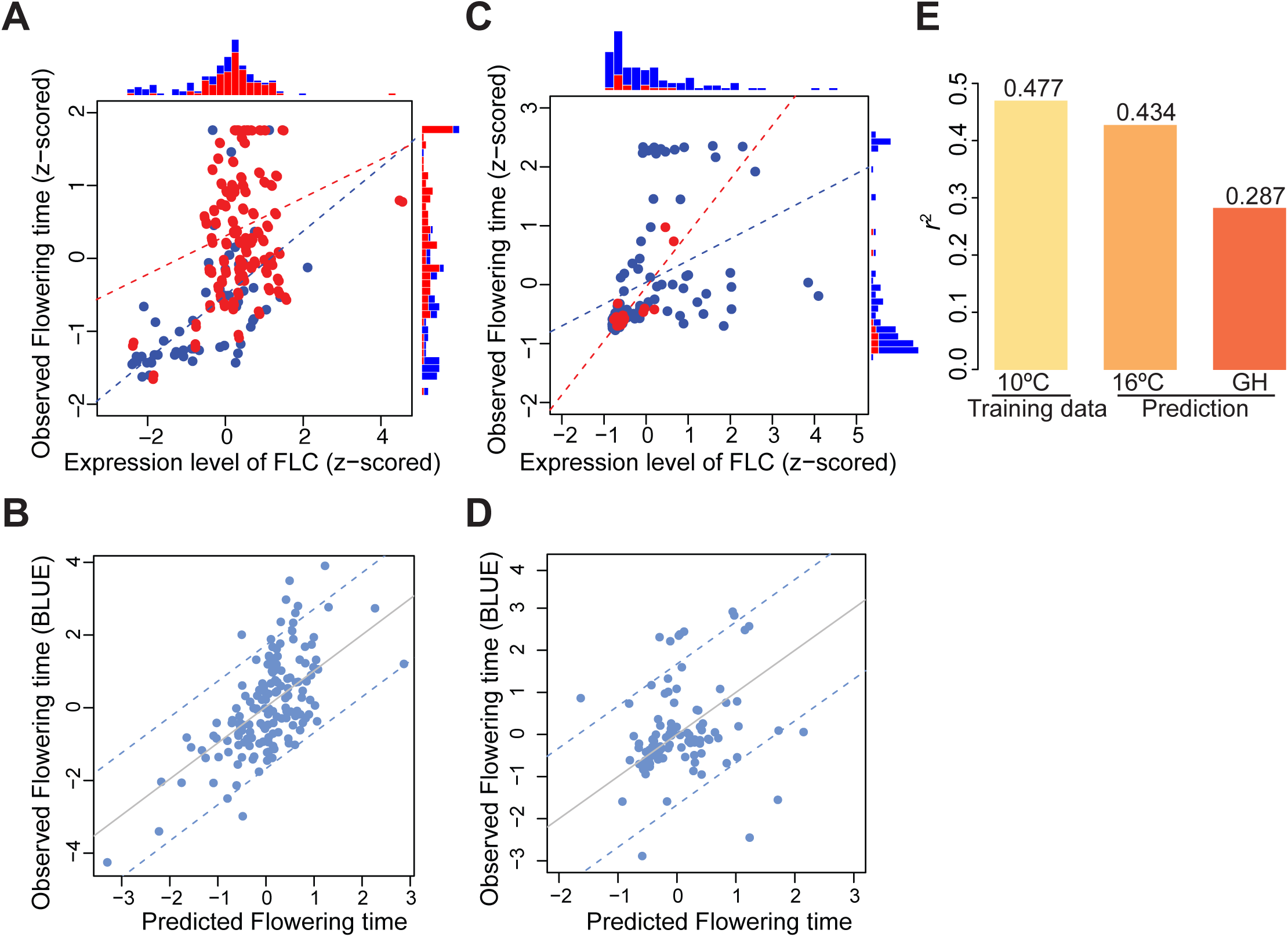
Prediction of flowering time by the *FLC*_10*°* *C*_ model. The data sets of the Swedish population (n=153; A, B) and global populations (n=101; C, D) were collected under 16*°*C and an ambient temperature in a greenhouse, respectively. (A, C) Scatter plots between flowering time and the expression level of *FLC*, and histograms showing distributions of the genotype SNP_*FLC*_. Blue and red show reference and non-reference allele, respectively. Dot lines showed regression lines for each allele. (B, D) Scatter plots between observed and predicted flowering time by the *FLC*_10_*°* _*C*_ full model. Dot lines show 95% confidence intervals. (E) Prediction of flowering time variation.

## Discussion

Our primary goal in this study was to explore how we might use transcriptome data to elucidate the genetic architecture and the regulatory network of a complex adaptive trait. Through integration analysis, we identified an extremely simple network structure that is determining flowering time in our condition (constant 10*°*C growth temperature in long day). Before discussing this in detail, it is worth noting that our overall results are very different from “typical” GWAS results in at least two ways. First, we find large allelic effects, and there is little “missing heritability” (MANOLIO *et al.* 2009) — the genetic variance explained by kinship alone (the “SNP heritability”) is consistent with direct estimates of heritability derived by comparing within and between line variances (ATWELL *et al.* 2010). Using a variance component approach (SASAKI *et al.* 2015), we estimated that alleles of the major flowering regulator *FLC* jointly explain 30% of the flowering time variation at 10*°*C, with the rest of the genome accounting for 56%. The existence of a major allelic variation is similar to what has been seen for some other locally adaptive traits, e.g., skin and eye color in humans (BELEZA *et al.* 2013), and is readily explained by selection maintaining variation. The high SNP heritability is presumably due to a combination of low environmental noise and high linkage disequilibrium leading to efficient capture of background genetic effects.

Second, SNPs detected in our GWAS are massively overrepresented in experimentally verified regulatory pathways directly related to flowering (Figure 3; SASAKI *et al.* 2015). This is very unlike most human traits, which mostly seem to vary due to pleiotropic mutations across the genome (BOYLE *et al.* 2017), but more similar variation in adaptively varying traits like skin and eye color (BELEZA *et al.* 2013). This agrees with the simple evolutionary expectation that adaptive variation should be less pleiotropic, whereas variation that is due to mutation-selection balance can affect any gene.

Indeed, not only are the GWAS hits directly related to flowering time, but the expression level associations are as well. (in agreement with several previous *A. thaliana* studies, e.g., SUBRAMANIAN *et al.* (2005) and JIMENEZ-GOMEZ *et al.* (2010). Using correlation between flowering time and transcriptome, we identified a gene list with a strong overrepresentation of known candidates (Table 1). Interestingly, with striking exception of *FLC*, there is no overlap between this list and the list of candidates identified by GWAS (SASAKI *et al.* 2015), suggesting that most of the genes on the former list are responding to genes on the latter list. This is certainly true for the small cluster of *FT* and *SOC1* under negative regulation by *FLC* (Figure 1B; KIM *et al.* 2009; WELLMER and RIECHMANN 2010). While the expression of all three genes is strongly correlated with flowering (and have been used as markers, *e.g.*, SATAKE *et al.* 2013; WANG *et al.* 2014; LEAL VALENTIM *et al.* 2015), only *FLC* appears to be directly causative, at least under this experimental condition. It is also notable that, with the obvious exception of *FLC*, genes that do harbor causative genetic variation do not show up as correlated in expression (Table 1). For example, expression levels of *VIN3*, a classical expression marker used in modeling (SATAKE *et al.* 2013), are not correlated with flowering despite *VIN3* having an apparent genetic effect (Fig 3A-B). Studies have shown that *VIN3* expression gradually increases during cold exposure, and that the abundance after sufficient long periods of exposure does not affect flowering time (WOLLENBERG and AMASINO 2012). Thus the reason for the lack of correlation in our study could be that the expression of *VIN3* was already saturated at this developmental stage (alternatively, genetic variation at *VIN3* could act at the amino-acid level).

Our analysis confirms that *FLC* plays a major role in determining flowering behavior (SHINDO *et al.* 2005; LI *et al.* 2014a), both in terms of being directly causative, and in terms of integrating variation at other loci. Importantly, *FLC* remains difficult to identify using standard, single-SNP, GWAS methods, the reasons being the complex genetic architecture of the locus itself. The situation is similar to that for the multi-allelic flowering locus *FRIGIDA* (SHINDO *et al.* 2005; ATWELL *et al.* 2010), but apparently much more complex (LI *et al.* 2014a). While SNP_*FLC*_ alone explained 19% of the phenotypic variation, local genetic variation at *FLC* explains 28%, and our full *FLC* model (including some *trans*-effects mediated by *FLC*) explains close to 50%. It is also notable that our estimate of the amount of the heritability that is attributable to expression is again much higher than in human disease studies. (O'CONNOR *et al.* 2017). It may seem paradoxical that our model, parametrized at 10*°*C, also works well at 16*°*C — and even at 23*°*C in a different population where the *cis*-regulatory variation at *FLC* is different, whereas the list of genes correlating with flowering time changed greatly between 10*°*C and 16*°*C (Table S3). The reason for this is not entirely clear, but likely involves the strong and locally adapted genetic background (LI *et al.* 2014a) which, to a significant extent, acts through *FLC* (as genotype-environment interactions). In conclusion, our novel mediation analysis illustrates the complexity of the genotype-phenotype map in even an extremely simple network dominated by a single locus (Figure 4A), but raises hope for more mechanistic (and genuinely predictive) models of the flowering time network (*e.g.*, ANGEL *et al.* 2015).

## Materials and Methods

### Correlation analysis

Data sets of 132 Swedish lines grown under constant 10*°*C were used for the analysis (LONG *et al.* 2013); DUBIN *et al.* (2015); SASAKI *et al.* (2015); Table S1). Correlation coefficient (*ρ*) was calculated between flowering time and expression levels for 20,285 genes for which more than 10% lines showed detectable expression levels. Also *r*^2^ and *p*-value were calculated by a general linear regression model using lm() function in R (www.r-project.org). Next, we calculated rho and the *p*-values for all pairs of gene and flowering time in Table 1. Using the significance, a correlation network was visualized using Cytoscape (SHANNON *et al.* 2003) with threshold *p*-value < 0.01 with bonferroni correction (741 tests for 38 genes + flowering time).

### GO analysis

Enrichment of known flowering time genes was estimated using BiNGO as a plugin of Cytoscape (MAERE *et al.* 2005), and Benjamini and Hochberg False Discovery Rate correction (BEN-JAMINI 1995) was used for the multiple testing correction. GO as “regulation of flower development” defined in the latest GO term in the Arabidopsis Information Resource (TAIR; BERARDINI *et al.* 2015) was used for the analysis as flowering time genes. FDRwas calculated based on the GO list as described in SASAKI *et al.* (2015).

### Linear mixed model (LMM)

All association studies were performed using LIMIX (LIPPERT *et al.* 2014). The following linear mixed model (LMM) was used

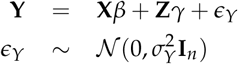

where **X** is the genotype of the SNP_*FLC*_ and *β* is the parameter of the corresponding fixed effect, **Z** = (*X*_1_ … *X*_*p*_) is all other SNPs and *γ ∼ N*(0, *σ*^2^ **I**_*p*_) is the corresponding random vector modeling the genomic background (KANG *et al.* 2008). Finally *I*_*n*_ is the *n* × *n* identity matrix.

To study the effect of gene expression with correction for population structure, the following LMM was used

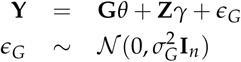

where **G** is the gene expression level and *θ* is the parameter for the corresponding fixed effect.

### Variance component analysis

*Cis*-genetic effects of loci on an expression level **Y** was estimated using local_vs_global_mm() function in mixmogam (https://github.com/bvilhjal/mixmogam) with the model

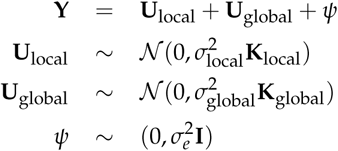

Where **U**_local_ and **U**_global_ are random effects corresponding to local and global relatedness, respectively, and *ψ* is noise. The local region is defined as +/- 15Kbp coding region of each gene. Significance of the variance component was estimated by permutation tests (1000 times) with maintaining the chromosomal order of all observations but shuffling the relative positions of the two variables (Figure S3).

### Mediation analysis

SAS macros published in VALERI and VANDERWEELE (2013) were used for this analysis. We implemented a linear mixed model to correct population structure to the model described in Supplemental Note. *r*^2^ was calculated as described in NAKAGAWA and SCHIELZETH (2013).

### Prediction of flowering time

Data sets for prediction are published flowering time and *FLC* expression data that were collected under constant 16*°*C growth temperature (DUBIN *et al.* 2015; SASAKI *et al.* 2015) and an ambient temperature around 23*°*C in a greenhouse (SHINDO *et al.* 2005; ATWELL *et al.* 2010). Lines included in the Swedish genome project (LONG *et al.* 2013) and the 1001 project (THE 1001 GENOMES CONSORTIUM 2016) were used for the analysis with the genotype (16*°*C n=153; greenhouse n=101; Table S1). The following model parameterized by the 10*°*C data set was used for prediction of flowering time **Y** including n individuals.

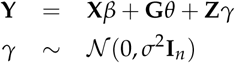

where **X** is the genotype of SNP_*FLC*_ and **G** is the expression levels of *FLC* under each condition. Based on the assumption that effects of population structure on **Y** and **G** are proportional, we estimate *γ* by fitting a null model **G** = **Z***γ* + *ψ* by REML implemented in EMMA (KANG *et al.* 2008). Flowering time variation explained by the model was estimated by *r*^2^ of NAKAGAWA and SCHIELZETH (2013). Significance of the variance component was estimated by permutation tests (1000 times) with maintaining the chromosomal order of all observations but shuffling the relative positions of the two variables (Figure S3).

